# AlphaFold accurately predicts distinct conformations based on oligomeric state of a de novo designed protein

**DOI:** 10.1101/2022.02.02.478886

**Authors:** Matthew C. Cummins, Tim M. Jacobs, Frank D. Teets, Frank DiMaio, Ashutosh Tripathy, Brian Kuhlman

**Affiliations:** Department of Pharmacology, University of North Carolina School of Medicine, Chapel Hill, North Carolina 27599; Department of Bioinformatics and Computational Biology, University of North Carolina School of Medicine, Chapel Hill, North Carolina 27599; Department of Biochemistry, University of Washington, Seattle, WA; Department of Biochemistry and Biophysics, University of North Carolina School of Medicine, Chapel Hill, North Carolina 27599; Lineburger Comprehensive Cancer Center, University of North Carolina at Chapel Hill, Chapel Hill, North Carolina 27599

**Keywords:** AlphaFold, machine learning, structure prediction, Rosetta, *de novo* design, G proteins

## Abstract

Using the molecular modeling program Rosetta, we designed a de novo protein, called SEWN0.1, that binds the heterotrimeric G protein Gαq. The design is helical, well-folded, and primarily monomeric in solution at a concentration of 10 uM. However, when we solved the crystal structure of SEWN0.1, we observed a dimer in a conformation incompatible with binding Gαq. Unintentionally, we had designed a protein that adopts alternate conformations depending on its oligomeric state. Recently, there has been tremendous progress in the field of protein structure prediction as new methods in artificial intelligence have been used to predict structures with high accuracy. We were curious if the structure prediction method AlphaFold could predict the structure of SEWN0.1 and if the prediction depended on oligomeric state. When AlphaFold was used to predict the structure of monomeric SEWN0.1, it produced a model that resembles the Rosetta design model and is compatible with binding Gαq, but when used to predict the structure of a dimer, it predicted a conformation that closely resembles the SEWN0.1 crystal structure. AlphaFold’s ability to predict multiple conformations for a single protein sequence should be useful for engineering protein switches.

## INTRODUCTION

DeepMind’s AlphaFold is a revolutionary method for protein structure prediction that has demonstrated high accuracy in community-wide assessments^1^ and has been used to predict the structures of 98.5% of the human proteome^2^. One exciting finding was that even though AlphaFold was trained to predict the structures of single protein chains, it can also accurately predict the structures of homo-oligomers ^1,3^. Since then, DeepMind has released AlphaFold-Multimer, a version of the program that is explicitly parameterized to predict the structures of protein homo- and hetero-complexes ^4^. Additionally, it has been demonstrated that AlphaFold can also be used for protein design. AlphaDesign is a new method for protein design that uses confidence metrics from AlphaFold as score terms during the in silico evolution of de novo sequences. Sequences designed with AlphaDesign have not been experimentally validated yet, but the method produces a large variety of sequences predicted with high confidence by AlphaFold to adopt specific three-dimensional structures. One interesting finding was that AlphaDesign can be used to design protein sequences that AlphaFold predicts to adopt alternative conformations based on oligomerization state, a behavior that has been observed in naturally occurring proteins^5,6^. This result suggests that AlphaFold may be a valuable tool for designing protein switches. Here, we add experimental support for this hypothesis as we show that AlphaFold accurately predicts an alternative conformation for a de novo designed protein that unintentionally crystallized as a dimer.

Our design goal was to create a small protein that binds and inhibits the heterotrimeric G protein Gα_q_. Gα_q_ is an oncogene that drives almost 90% of uveal melanomas^7,8^. We previously reported that polypeptides that compete with the binding of Gα_q_’s native effector, PLCβ3, inhibit Gα_q_ signaling^9^. To design additional competitive inhibitors, we used the SEWING design protocol in Rosetta to embed a helix-turn-helix (HTH) motif from PLCβ3^10^ (Fig. 1A) in a de novo designed protein. This strategy was employed because the HTH motif makes extensive contacts with Gα_q_, and peptides based on the HTH can inhibit Gα_q_ signaling. The SEWING protocol constructs novel protein backbones by piecing together small fragments of naturally occurring proteins^11,12^. In this case, the HTH motif was included as a starting building block. The last step of the design protocol is rotamer-based sequence optimization in Rosetta^13–15^. From thousands of design models, we picked three sequences for experimental validation. Here, we describe the crystal structure of one of these sequences and compare it with the Rosetta model and models generated with AlphaFold. AlphaFold was not available when the sequences were first designed.

**Figure 1.**
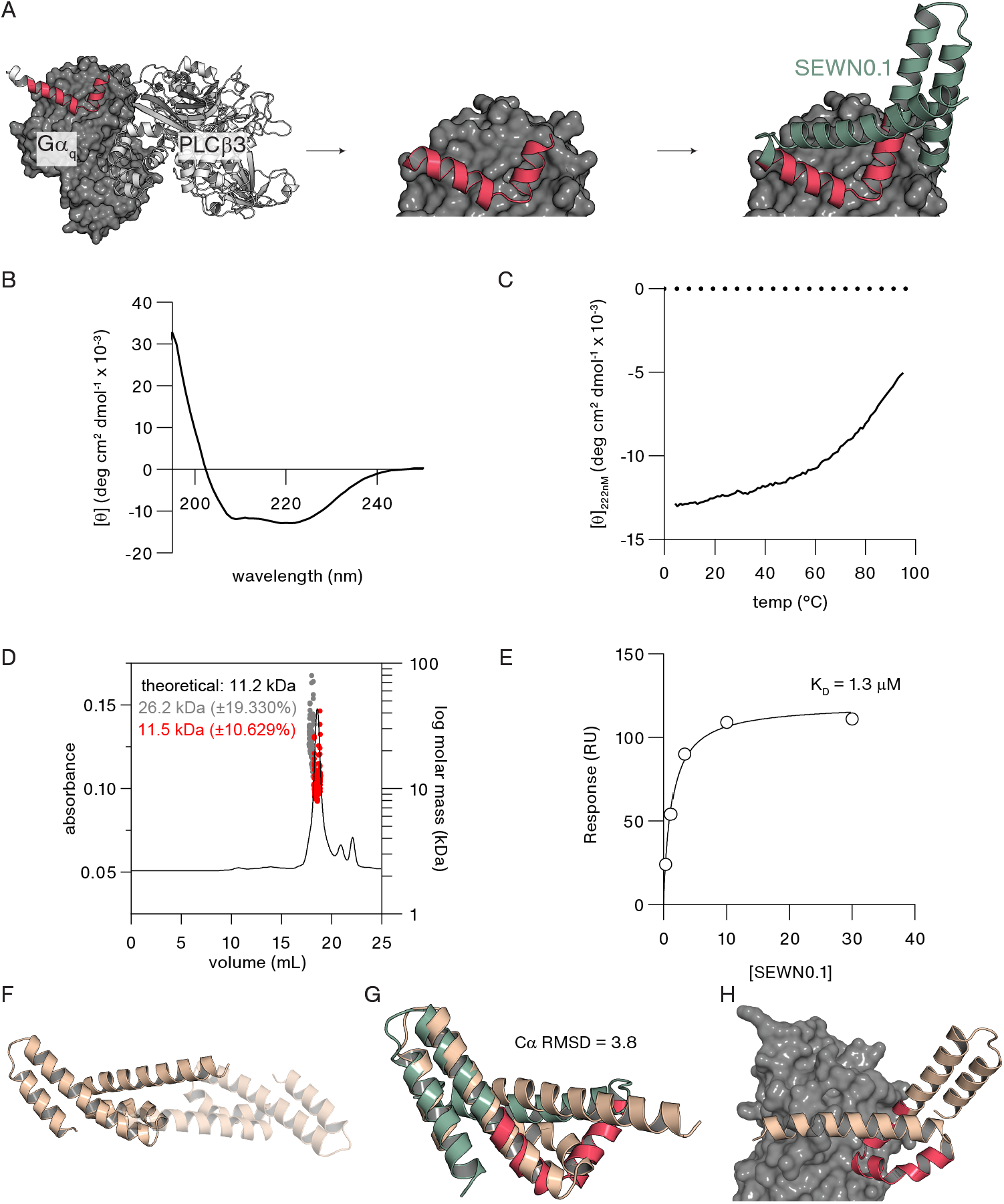
**A.** To create a G*α_q_* inhibitor, residues 853-876 (magenta) from PLCβ3 (PDB 7SQ2) were embedded in a de novo designed protein using the SEWING protocol in Rosetta. **B.** CD spectra of SEWN0.1. **C.** Thermal denaturation of SEWN0.1 as monitored by CD signal at 222nm. **D.** SEC-MALS experiment with 10 μM SEWN0.1 demonstrates that the protein is primarily monomeric in solution **E.** SEWN0.1 binding biotinylated Gα_q/I_ as monitored with SPR. **F.** In the crystal structure of SEWN0.1 the protein forms a dimer and **G.** adopts a different conformation than the design model (the design model is shown in green with the helix-turn-helix motif in magenta). **H.** The alternative conformation is incompatible with binding to Gα_q_ via the helix-turn-helix motif (magenta) as it directs helix 4 into the surface of the protein.

## RESULTS AND DISCUSSION

### SEWN0.1 is a monomeric, stable, and helical protein that binds Gα_q/i_ at the desired paratope

The SEWING protocol was used to design three all-helical proteins that incorporate the HTH motif from PLCβ3. During assembly of the de novo backbones, protein conformations were disallowed that would result in steric clashes when the HTH engages Gα_q_. During sequence design, amino acids from the HTH that directly interact with Gα_q_ were not allowed to mutate, but residues on the backside of the motif that do not interact with Gα_q_ were mutated to make favorable contacts with the rest of the de novo protein. The three proteins were expressed and purified from *E. coli*, and we were able to solve the crystal structure of one of them, named SEWN0.1. SEWN0.1 is a 93-residue protein predicted to have four α-helices. A BLAST search of the non-redundant protein sequences database (NCBI) produced zero hits with an E-value below 0.5.

SEWN0.1 has a circular dichroism (CD) spectrum indicative of a helical protein (Fig. 1B) and unfolds cooperatively with temperature (Fig. 1C). Size exclusion chromatography coupled with multi-angle light scattering confirmed that the protein is predominantly monomeric at a concentration of 10 uM (Fig. 1D). The molecular weight (MW) was measured to be 11.5 kDa (±10.6%), close to the theoretical MW of 11.2 kDa. However, the scattering data also indicated the presence of larger molecular weight species suggesting that the protein may have the propensity to homooligomerize. The equilibrium dissociation constant (K_D_) between SEWN0.1 and Gα_q_ was measured by surface plasmon resonance to be 1.3 ± 0.1 μM (Fig. 1E, Fig. S2). To determine if SEWN0.1 binds to the target binding site on Gα_q_ we performed a competitive fluorescence polarization experiment with a fluorescently labeled HTH-peptide derived from PLCβ3. As SEWN0.1 was titrated into a solution of HTH-peptide and Gα_q_ we observed the fluorescence polarization signal decrease, confirming that SEWN0.1 binds to the target binding site (Fig. S3).

### SEWN0.1 crystallizes as a dimer in an alternative conformation

We solved the crystal structure of SEWN0.1 to a resolution of 1.9 Å (Table S1). To our surprise, SEWN0.1 crystallized as a dimer (Fig. 1F) in a different conformation from the Rosetta model, Cα RMSD = 3.8Å (Fig. 1G). As was intended, SEWN0.1 consists of four α-helices, but the third (Hα3) and fourth (Hα4) helices point in opposite directions in the crystal structure than in the design model. The alternative conformation results in an exposed hydrophobic surface on Hα4 that is buried at the dimer interface. One of the most striking differences between the model and crystal structure is the loop that connects Hα2 with Hα3. This loop is meant to mimic the HTH motif from PLCβ3. Furthermore, the conformation in the crystal structure is not compatible with Gαq binding. When SEWN0.1 is superimposed onto the crystal structure of the PLCβ3/Gαq complex (by superimposing Hα2 and Hα3 onto the HTH motif), Hα4 drives straight into Gαq (Fig. 1H). Lastly, the buried surface area of the homodimer is ~2100Å, or 20% of the solvent-accessible surface area, indicating that it is likely to be more than just a weak crystal contact^16^. Taken together, our results suggest that SEWN0.1 adopts a different conformation in the crystal than when it is a soluble monomer at micromolar concentrations.

### AlphaFold predicts alternative conformations for monomeric and dimeric SEWN0.1

AlphaFold has been shown to predict the structures of monomers^1^ and complexes^4^ with high accuracy. Since we have evidence that SEWN0.1 adopts alternative conformations depending on oligomeric state, we reasoned that it would be an exciting test for AlphaFold. Impressively, AlphaFold predicts two distinct conformations for SEWN0.1 depending on oligomeric state. As a monomer, AlphaFold predicts SEWN0.1 to adopt a very similar structure to the Rosetta model, Cα RMSD = 0.82 (Fig. 2A) and is consistent with binding to Gα_q_. However, as a dimer, AlphaFold predicts SEWN0.1 to adopt a conformation that resembles the crystal structure, Cα RMSD = 1.9 (Fig. 2B). In the dimer model, the Hα3 and Hα4 helices point in the same direction as the crystal structure. Interestingly, AlphaFold did not accurately predict the relative orientation of the monomers at the dimer interface (Fig. 2C). This result could be because of crystal packing effects. The AlphaFold prediction of the dimer was also sensitive to the precise sequence used for the N-terminal region of the protein. If the sequence resolved in the SEWN0.1 crystal structure (starts at residue 6) is used for the AlphaFold prediction, then the prediction matches the crystal structure as shown in Figure 2. The same prediction is made if unresolved residues 4 and 5 are included. However, if residues 1-3 are included in the sequence, then the AlphaFold model reverts to the Rosetta design model. Given that these residues are not resolved in the electron density, it is not clear why they influence the AlphaFold prediction, but the result does highlight the chameleon nature of SEWN0.1. This structural ambiguity is reflected in the confidence predictions output by AlphaFold. The loop connecting helices 2 and 3 has the largest predicted error in the AlphaFold results for both the monomer and the dimer (Fig. S4).

**Figure 2.**
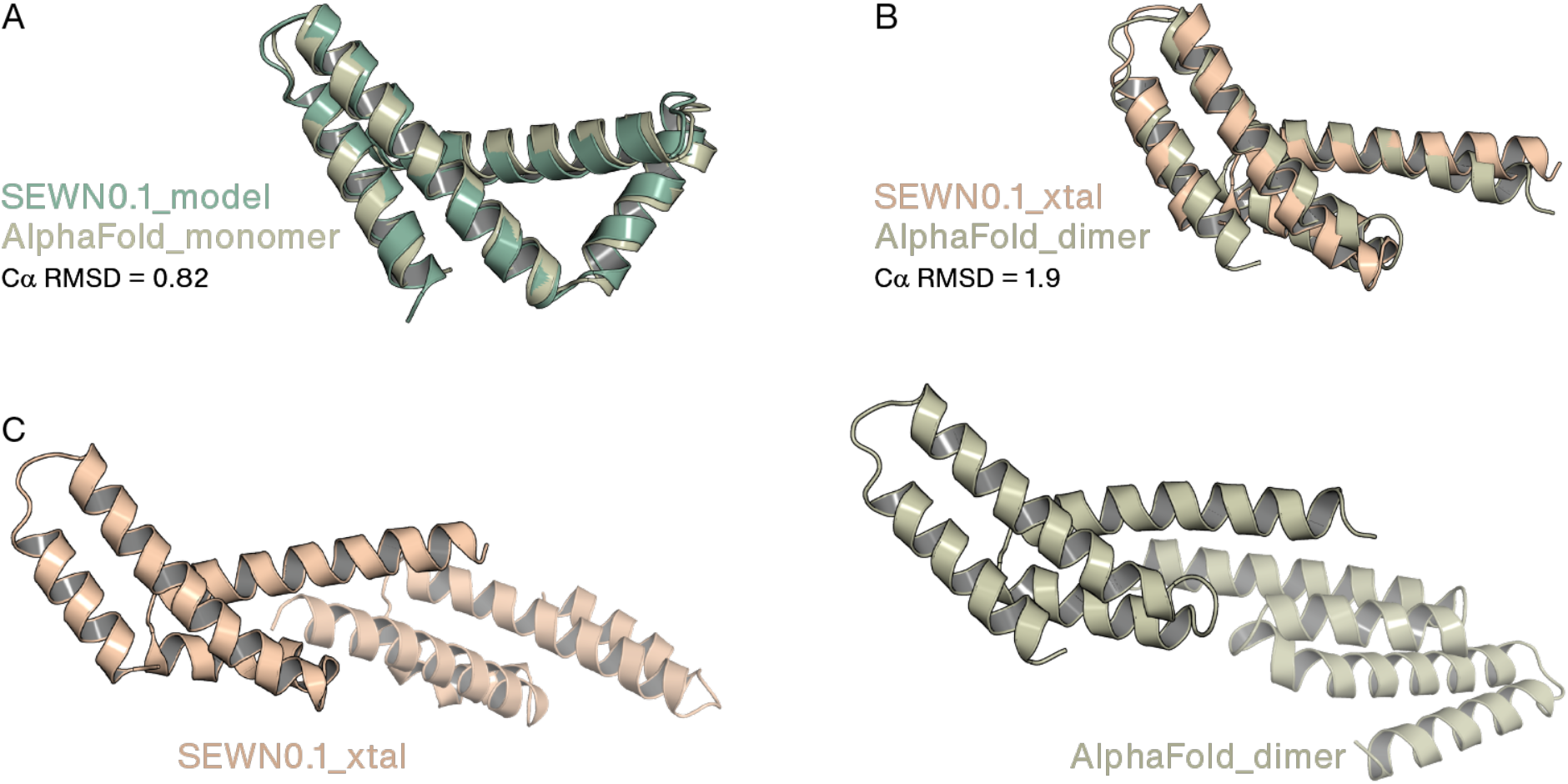
AlphaFold accurately predicts distinct conformations based on oligomeric state. **A.** Superimposition of SEWN0.1 Rosetta model onto AlphaFold monomer prediction. **B.** Superimposition of one protomer of SEWN0.1 crystal structure and one protomer of AlphaFold dimer prediction. **C.** AlphaFold does not accurately predict the dimer interface

## CONCLUSION

The structure prediction of SEWN0.1 is a rigorous test for AlphaFold. It is a de novo sequence with no natural homologs, and as an unpublished structure, it was not included in the training of AlphaFold. Furthermore, we demonstrated that the structure of SEWN0.1 is dependent on oligomerization state. The ability of AlphaFold to accurately predict the conformation of the SEWN0.1 dimer highlights the capabilities of AlphaFold and suggests that it will be a valuable tool for the design of proteins that switch conformation upon oligomerization. Indeed, the AlphaDesign method has used AlphaFold to design sequences predicted to switch conformation upon oligomerization; however, these predictions have not been experimentally verified yet^17,18^. It is interesting to consider why the AlphaFold prediction for SEWN0.1 changes when forced to model a homodimer. In training, AlphaFold has almost certainly learned to maximize the burial of hydrophobic surface area. The formation of a homodimer creates an additional mechanism for burying residues. It appears that AlphaFold detected that a conformational change in the loop between helix 2 and 3 would expose hydrophobic residues on helices 3 and 4 that could then be packed at the dimer interface. In the future, it will be exciting to see if AlphaFold can be used to design heterocomplexes that induce conformational change predictably.

## MATERIALS AND METHODS

### Computational design of SEWING proteins

The SEWING protocol from a previous study^12^ was used with a few differences. Residues 853-876 from PLCβ3 (PDB: 3OHM) were used as a starting node for the SEWING assembly. Furthermore, Gα_q_ (PDB: 3OHM) was present to prevent the addition of segments that would sterically clash with Gα_q_ if bound at the desired paratope. The sequences of the helical assemblies were optimized using iterative rounds of rotamer-based repacking and energy-based minimization of torsion angles^13–15^. Finally, the three top-scoring designs were tested for experimental validation.

### Recombinant expression and purification

Expression and purification of SEWING proteins and biotinylated Gα_q/I_ were performed using the same methods described previously^9^. Briefly, after recombinant expression in *E. coli* cells, proteins were purified using Ni-NTA IMAC. After cleaving and removing the His-tag, protein samples were purified using an S75 and GE AKTA FPLC system. With Gα_q/i_, when the protein was bound to Ni-NTA resin, the sample had an additional incubation step with BirA and Biomix B (Avidity) to install biotinylation.

### Circular Dichroism

CD experiments were performed using a Jasco J-815 CD spectrometer. Samples were prepared in 25 mM sodium phosphate pH 7.0 and 50 mM NaCl at a concentration of 30 μM. Measurements were collected at 20°C, at scanning speed of 50 nm/min, 2-second DIT, and 2 nm bandwidth. Thermal denaturation was performed by ramping the temperature 1°C/min from 4°C to 95°C and measuring the CD signal at 222 nm.

### SPR

Biotinylated Gα_q/i_ was loaded onto a neutravidin sensor chip (ProteOn NLC) at a concentration of 100 nM. The three SEWING proteins, SEWN0.1-3, flowed over the chip at 5 concentrations from 0.37 μM to 30 μM. The collected data fit a single-state equilibrium binding model. All experiments were performed in 20 mM HEPES pH 7.0, 10 mM NaF, 30 μM AlCl_3_, 10 mM MgCl_2_, 150 mM NaCl, 2 mM DTT, and 0.005% Tween-20.

### SEC-MALS

SEC-MALS experiments were performed using a Wyatt DAWN HELEOS II light scattering instrument connected to an Agilent FPLC System and Wyatt T-rEX refractometer. The sample was run over a Superdex S200 10/300 column at 0.5 mL/min flowrate. The system was equilibrated in 25mM sodium phosphate, 250mM NaCl, 0.05% sodium azide, pH 7.0. The protein sample concentration was 20 μM. The experiment was performed at room temperature. The data was analyzed using Wyatt ASTRA 6.

### Crystallography and structure determination

Purified SEWN0.1 was concentrated to 225 μM and dispensed in 24-well crystallography trays (Hampton Research). The sample crystallized in hanging drops via vapor diffusion in 100 mM diammonium hydrogen citrate and 15% polyethylene glycol after approximately 3 days. Crystals were soaked in a mother liquor solution mixed with 30% ethylene glycol before being flash-frozen in liquid nitrogen. Crystals were shot at the Advanced Photon Source, Argonne National Laboratory using the Southeast Regional Collaborative Access Team 22-ID beamline. Data were processed and scaled with XDS^19^. Initial phases were solved by molecular replacement using PHASER^20^ and residues 6-43 from the Rosetta design model. SHELXE^21^ was used to build a poly-alanine backbone that was built and refined using COOT^22^ and Phenix-Refine^23^. Buried surface area was calculated using Areaimol from the CCP4 suite^24^. Coordinates were deposited to the PDB (PDB: 7TJL).

### Competitive fluorescence anisotropy

Competitive FA assays were performed the same as in a previous study^9^. Briefly, 400 nM of TAMRA-labeled HTH-pep was incubated with 2 μM of Gα_q/i_. Fluorescence anisotropy was measured upon increasing titration of SEWN0.1 at the indicated concentrations. The experiment was performed on a Jobin Yvon Horiba FluoroMax3 fluorescence spectrometer in 20 mM HEPES pH 7.4, 150 mM NaCl, 1 mM MgCl2, 30 μM AlCl3, and 10 mM NaF.

### AlphaFold predictions

AlphaFold predictions were made using the AlphaFold2.ipynb v1.0 colab notebook as part of the ColabFold framework^3^. The predictions did not use a multiple sequence alignment (msa_mode = single sequence) and were tested using homooligomer 1 (monomer) or 2 (homodimer). DeepMind released 5 different weight sets for AlphaFold. However, choice of weight set did not dramatically change the structure predictions for the monomer or dimer.

## ACKNOWLEDGMENTS

This work was supported by NIH grant R35GM131923(BK) and T32GM007040(MC).

## References

1. Jumper J, Evans R, Pritzel A, Green T, Figurnov M, Ronneberger O, Tunyasuvunakool K, Bates R, Žídek A, Potapenko A, et al. (2021) Highly accurate protein structure prediction with AlphaFold. Nature 596:583–589.

2. Tunyasuvunakool K, Adler J, Wu Z, Green T, Zielinski M, Žídek A, Bridgland A, Cowie A, Meyer C, Laydon A, et al. (2021) Highly accurate protein structure prediction for the human proteome. Nature 596:590–596.

3. Mirdita M, Ovchinnikov S, Steinegger M (2021) ColabFold - Making protein folding accessible to all. bioRxiv:2021.08.15.456425.

4. Evans R, O’Neill M, Pritzel A, Antropova N, Senior A, Green T, Žídek A, Bates R, Blackwell S, Yim J, et al. (2021) Protein complex prediction with AlphaFold-Multimer. bioRxiv:2021.10.04.463034.

5. Penke B, Szűcs M, Bogár F (2020) Oligomerization and Conformational Change Turn Monomeric β-Amyloid and Tau Proteins Toxic: Their Role in Alzheimer’s Pathogenesis. Molecules 25:1659.

6. Edelstein SJ, Changeux J-P (2010) Relationships between Structural Dynamics and Functional Kinetics in Oligomeric Membrane Receptors. Biophys. J. 98:2045–2052.

7. Van Raamsdonk CD, Griewank KG, Crosby MB, Garrido MC, Vemula S, Wiesner T, Obenauf AC, Wackernagel W, Green G, Bouvier N, et al. (2010) Mutations in GNA11 in Uveal Melanoma. N. Engl. J. Med. 363:2191–2199.

8. Van Raamsdonk CD, Bezrookove V, Green G, Bauer J, Gaugler L, O’Brien JM, Simpson EM, Barsh GS, Bastian BC (2009) Frequent somatic mutations of GNAQ in uveal melanoma and blue naevi. Nature 457:599–602.

9. Hussain M, Cummins MC, Endo-Streeter S, Sondek J, Kuhlman B (2021) Designer proteins that competitively inhibit Gαq by targeting its effector site. J. Biol. Chem. 297:101348.

10. Waldo GL, Ricks TK, Hicks SN, Cheever ML, Kawano T, Tsuboi K, Wang X, Montell C, Kozasa T, Sondek J, et al. (2010) Kinetic Scaffolding Mediated by a Phospholipase C–β and G q Signaling Complex. Science 330:974–980.

11. Guffy SL, Teets FD, Langlois MI, Kuhlman B (2018) Protocols for Requirement-Driven Protein Design in the Rosetta Modeling Program. J. Chem. Inf. Model. 58:895–901.

12. Jacobs TM, Williams B, Williams T, Xu X, Eletsky A, Federizon JF, Szyperski T, Kuhlman B (2016) Design of structurally distinct proteins using strategies inspired by evolution. Science 352:687–690.

13. Fleishman SJ, Leaver-Fay A, Corn JE, Strauch E-M, Khare SD, Koga N, Ashworth J, Murphy P, Richter F, Lemmon G, et al. (2011) RosettaScripts: A Scripting Language Interface to the Rosetta Macromolecular Modeling Suite Uversky VN, editor. PLoS One 6:e20161.

14. Khatib F, Cooper S, Tyka MD, Xu K, Makedon I, Popovic Z, Baker D, Players F (2011) Algorithm discovery by protein folding game players. Proc. Natl. Acad. Sci. 108:18949–18953.

15. Bhardwaj G, Mulligan VK, Bahl CD, Gilmore JM, Harvey PJ, Cheneval O, Buchko GW, Pulavarti SVSRK, Kaas Q, Eletsky A, et al. (2016) Accurate de novo design of hyperstable constrained peptides. Nature 538:329–335.

16. Elez K, Bonvin AMJJ, Vangone A (2018) Distinguishing crystallographic from biological interfaces in protein complexes: role of intermolecular contacts and energetics for classification. BMC Bioinformatics 19:438.

17. Jendrusch M, Korbel JO, Sadiq SK (2021) AlphaDesign: A de novo protein design framework based on AlphaFold. bioRxiv:2021.10.11.463937.

18. Akdel M, Pires DE V, Pardo EP, Jänes J, Zalevsky AO, Mészáros B, Bryant P, Good LL, Laskowski RA, Pozzati G, et al. (2021) A structural biology community assessment of AlphaFold 2 applications. bioRxiv:2021.09.26.461876.

19. Kabsch W (2010) XDS. Acta Crystallogr. Sect. D Biol. Crystallogr. 66:125–132.

20. McCoy AJ, Grosse-Kunstleve RW, Adams PD, Winn MD, Storoni LC, Read RJ (2007) Phaser crystallographic software. J. Appl. Crystallogr. 40:658–674.

21. Thorn A, Sheldrick GM (2013) Extending molecular-replacement solutions with SHELXE. Acta Crystallogr. Sect. D Biol. Crystallogr. 69:2251–2256.

22. Emsley P, Cowtan K (2004) Coot: model-building tools for molecular graphics. Acta Crystallogr. Sect. D Biol. Crystallogr. 60:2126–2132.

23. Afonine P V., Grosse-Kunstleve RW, Echols N, Headd JJ, Moriarty NW, Mustyakimov M, Terwilliger TC, Urzhumtsev A, Zwart PH, Adams PD (2012) Towards automated crystallographic structure refinement with phenix.refine. Acta Crystallogr. Sect. D Biol. Crystallogr. 68:352–367.

24. Collaborative Computational Project N 4 (1994) The CCP4 suite: programs for protein crystallography. Acta Crystallogr. Sect. D Biol. Crystallogr. 50:760–763.

