## Supplementary Figures for "AlphaFold accurately predicts distinct conformations based on oligomeric state of a de novo designed protein"

Tim M. Jacobs, AbCellera Biologics Inc., 2215 Yukon St, Vancouver, BC V5Y 0A1

Frank D. Teets, Department of Computational Biology, AI Proteins, 4 Corporate Drive, Andover, MA 01810

**Table S1 Crystallography data collection and refinement statistics for SEWN0.1**

| **Data Collection** | |
|---|---|
| Wavelength | 1.0 |
| Resolution range | 100 - 1.87 (1.92 - 1.87) |
| Space group | C 1 2 1 |
| Unit cell | 116.25, 31, 31.36<br>90, 93.071, 90 |
| Total reflections | 9846 |
| Unique reflections | 9100 (455) |
| Completeness (%) | 93.30 (66.10) |
| Mean I/σ (I) | 15.48 (3.20) |
| Wilson B-factor | 21.08 |
| R-merge | 0.0550 (0.2440) |
| R-meas | 0.0650 (0.3260) |
| CC1/2 | 0.9960 (0.9210) |
| **Refinement** | |
| Reflections used in refinement | 8796 (607) |
| Reflections used for R-free | 882 (55) |
| R-work | 0.2103 (0.2722) |
| R-free | 0.2304 (0.3632) |
| Number of non-hydrogen atoms | 825 |
| macromolecules | 732 |
| ligands | 0 |
| solvent | 93 |
| Protein residues | 85 |
| RMS(bonds) | 0.009 |
| RMS(angles) | 1.01 |
| Ramachandran favored (%) | 100 |
| Ramachandran allowed (%) | 0 |
| Ramachandran outliers (%) | 0 |
| Rotamer outliers (%) | 0 |
| Clashscore | 2.74 |
| Average B-factor | 26.88 |
| macromolecules | 26.07 |
| solvent | 33.29 |

Note: values in parenthesis correspond to highest resolution bin

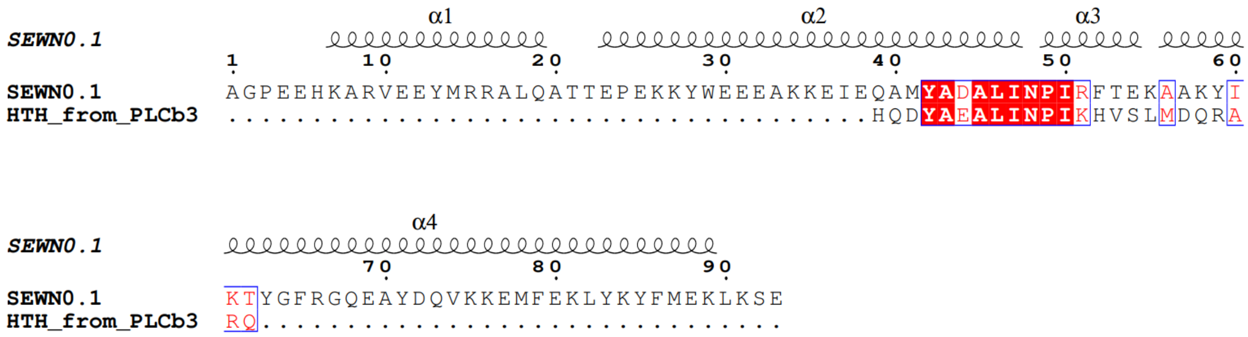


**Figure S1** Full sequence of SEWN0.1 aligned with HTH from PLCβ3
Secondary structure depiction is based on structure of SEWN0.1 (PDB 7TJL). Image of alignment was generated using ESPript 3.0.

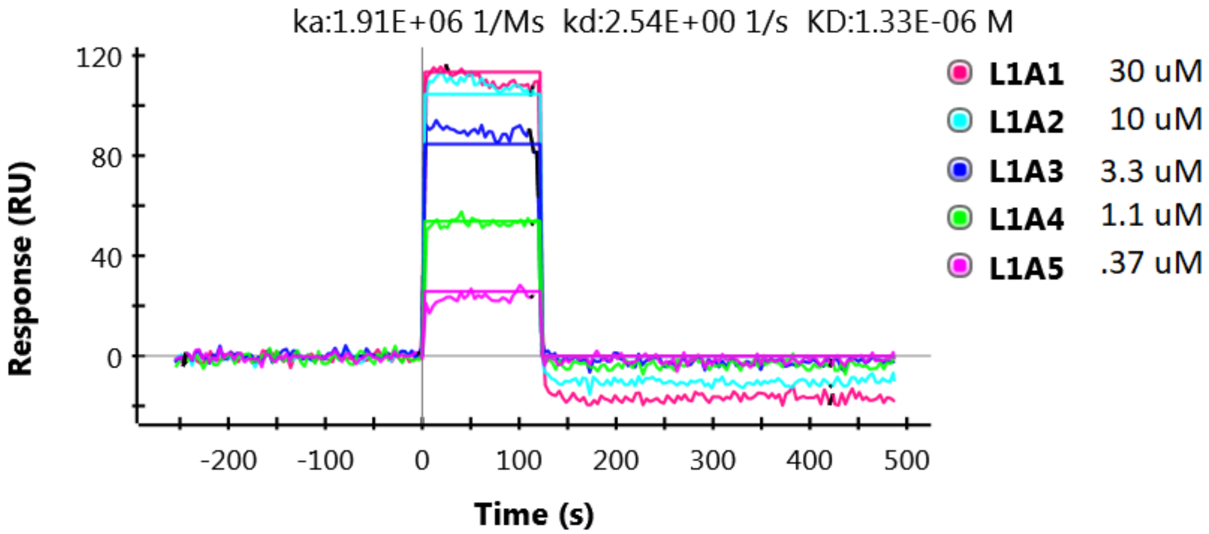


**Figure S2** Raw kinetic traces of SPR experiment of SEWN0.1 binding biotinylated G$\alpha_{q/i}$ Biotinylated G$\alpha_{q/i}$ was immobilized on a NeutrAvidin chip. SEWN0.1 was flowed over the chip at the 5 indicated concentrations. Binding affinity was determined by doing a kinetic analysis fit.

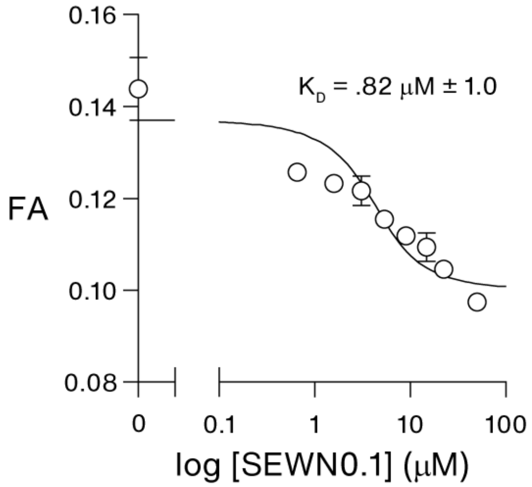


**Figure S3** Competitive fluorescence anisotropy experiment of SEWN0.1 binding $G\alpha_{q/i}$
$G\alpha_{q/i}$ was incubated with TAMRA-labelled HTH peptide. Fluorescence anisotropy was measured at the indicated concentrations of SEWN0.1. Data was numerically fit as described in Hussain, et al.

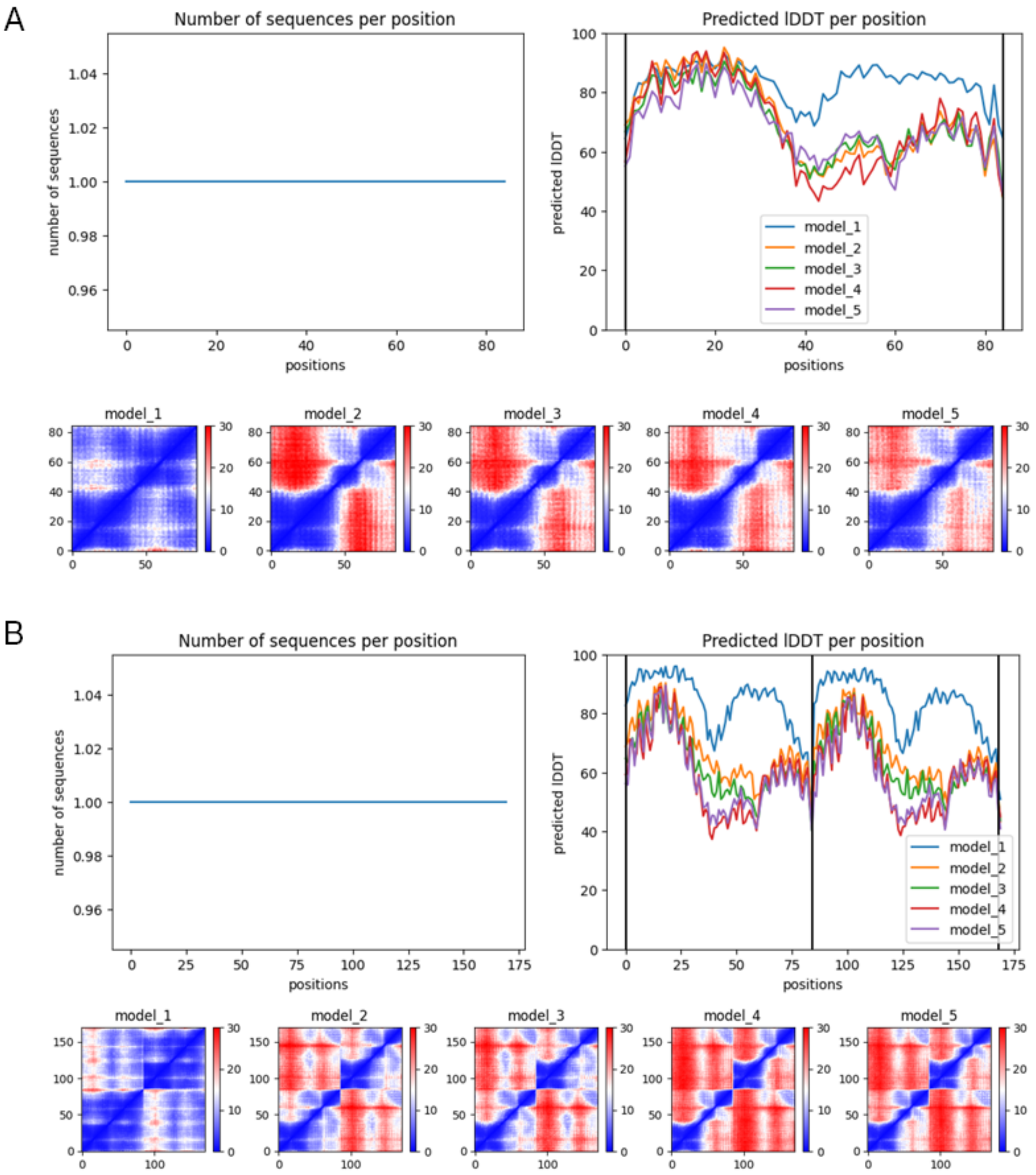


**Figure S4.** Predicted local Difference Distance Test (lDDT) and alignment errors graphs from AlphaFold prediction: **A.** monomer and **B.** dimer. Residue 43 corresponds to N48 in the full-length sequence and represents the single residue turn between Hα2 and Hα3. From the lDTT plots it is evident that AlphaFold is least confident in its prediction of the turn between Hα2 and Hα3, the region that adopts alternative conformations in the monomer and dimer models.
